# Autoturbo-DNA: Turbo-Autoencoders for the DNA data storage channel

**DOI:** 10.1101/2023.09.15.557887

**Authors:** Marius Welzel, Hagen Dreßler, Dominik Heider

## Abstract

DNA, with its high storage density and long-term stability, is a potential candidate for a next-generation storage device. The DNA data storage channel, comprised of synthesis, amplification, storage, and sequencing, exhibits error probabilities and error profiles specific to the components of the channel. Here, we present Autoturbo-DNA, a PyTorch framework for training error-correcting, overcomplete autoencoders specifically tailored for the DNA data storage channel. It allows training different architecture combinations and using a wide variety of channel component models for noise generation during training. It further supports training the encoder to generate DNA sequences that adhere to user-defined constraints.

## 1 Introduction

The exponential increase in data generation [1] leads to an increasing demand for data storage solutions with a high storage density and long-term stability. The global demand for data storage is estimated to reach 175 ZB in 2025 [2]. DNA, with a storage density of around 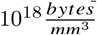, and a potential lifetime of hundreds of years [1], is one potential candidate for such a next-generation storage device. Utilizing DNA as a means to store data is an active field of research, with many advances that were reported in recent years [3, 4, 5, 6, 7, 8]. One important aspect of DNA as a data storage device is the error types that are typically observed in DNA: besides the change from one base to another (substitutions), other error types common in DNA are the insertion of a base into the DNA strand, leading to all bases that follow the erroneous base to be shifted one position to the right, or the deletion of a base, leading to all following bases being shifted one position to the left. The positional change of many bases at once increases the decoding complexity, and many different strategies have been developed for decoding sequences that contain indel (i.e., insertion or deletion) errors. Example strategies include encoding multiple copies of the same input block into multiple encoded blocks, which allows to compensate for some indel errors by falling back to other encoded blocks containing the same information as the erroneous block [9, 5]. Another approach presented in [10] is maximum likelihood tree decoding, utilizing periodic synchronization markers for decoding data in the presence of indel errors. A further approach is the exploitation of sequencing depth by using a form of majority voting for sequencing reads, which allows compensation for indels that occurred during sequencing [11]. Another important aspect is that the DNA data storage channel comprises multiple components, the writing of data into DNA (synthesis), the amplification of the synthesis product using PCR, the storage process itself, and the reading of the DNA back into a digital format (sequencing). Each component, including the various options for a component (e.g., different polymerases for the PCR or different sequencing machines), has specific error profiles and error patterns.^12^ Incorporation of this knowledge into the design process of coding schemes for DNA data storage could lead to more robust codes with less required redundancy. Furthermore, the encoded DNA sequences have to adhere to some specific constraints. The guanine and cytosine (GC) content in a sequence, compared to the adenine and thymine content, should be roughly balanced. Sequences with a GC content that deviates substantially from 50 % can lead to synthesis failures, unstable DNA, as well as sequencing errors. Similarly, large chains of the same base in a sequence (so-called homopolymers) can also lead to sequencing errors. The encoded DNA sequences should also be free of certain motifs specific to the synthesis or storage processes; for example, encoded sequences should not contain any restriction motifs used in the synthesis process or motifs with biological relevance if the encoded data is stored in-vivo. Further constraints, such as the occurrence of short repetitive sequences (k-mers) or the probability of secondary structure formation of the encoded sequences, must be also considered in some cases. One possible way to incorporate sequence constraints, error profiles, and error patterns into a coding system that can be flexibly adjusted to different combinations of synthesis, PCR, storage, and sequencing would be to leverage the learning ability of neural networks. Jiang et.al. [13] presented an end-to-end autoencoder coding system, TurboAE, that is solely based on neural networks (NN) but with a decoder that is arranged similarly to turbo codes [14]. The authors showed that for some non-canonical channels, TurboAE outperforms state-of-the-art codes. Given the complexity of the DNA data storage channel, NN-based coding systems like TurboAE can learn the peculiarities of the DNA storage channel while being able to be fine-tuned on specific setups of synthesis, sequencing, and storage methods and PCR polymerases, which could lead to improvements regarding the error correction performance and execution time required for the en- and decoding processes. Another potential advantage of such coding systems is that they perform well for short and moderate block lengths. For DNA, it is practical and efficient to have block lengths that correspond to strand lengths that can be synthesized and sequenced in one synthesis and sequencing process without requiring multiple reads to be assembled into one block during sequencing data processing. These block lengths are typically between 300 and 1000 base pairs for current sequencing technologies, while shorter fragment sizes are cheaper to synthesize on a dollar-per-base basis. Here, we present Autoturbo-DNA, an end-to-end autoencoder framework that combines the TurboAE principles with an additional pre-processing decoder, DNA data storage channel simulation, and constraint adherence check. Autoturbo-DNA supports various NN architectures for its components, which can be mixed and matched using a configuration file, combined with user-friendly adjustment of the DNA data storage channel and constraint adherence parameters.

## 2 Methods

### 2.1 Channel design

To train a model that can effectively repair erroneous sequences, the channel has to be modeled as close to the real DNA channel as possible. Autoturbo-DNA supports many different DNA data storage channel components, as shown in table 1. The supported error rates and patterns are based on literature data collected by [12]. Given the rapid development in DNA synthesis and sequencing, together with potential alternative storage media, like silica particles [15], error rates and -patterns need to be highly customizable. Autoturbo-DNA allows for such customization by utilizing JSON files for the channel parameters. The JSON files are cross-compatible with the configuration files used in the error simulator MESA [12]. This interoperability between Autoturbo-DNA and MESA allows users to use the MESA error probability customization GUI to generate complex error patterns using a simple-to-use, click-and-drag-based interface. An example of how to generate a JSON configuration file for the Autoturbo-DNA channel simulator using the MESA GUI is shown in the supplement.

**Table 1:** Out-of-the-box supported error rates for the channel simulation.

| Synthesis method | Error-correction | Deletion | Insertion | Substitution | Raw rate |
| --- | --- | --- | --- | --- | --- |
| Column synthesized oligos | ErrASE | 0.6 | 0.2 | 0.2 | $2.50 \cdot 10^{-5}$ |
| Microarray based oligo pools | ErrASE | 0.6 | 0.2 | 0.2 | $1.20 \cdot 10^{-3}$ |
| Column synthesized oligos | MutS | 0.7 | 0.15 | 0.15 | $1.00 \cdot 10^{-4}$ |
| Column synthesized oligos | Consensus Shuffle | 0.7 | 0.15 | 0.15 | $1.25 \cdot 10^{-4}$ |
| **PCR Polymerase** | **Proofread** |  |  |  |  |
| Taq | No | 0.01 | 0 | 0.99 | $4.30 \cdot 10^{-5}$ |
| Pwo | Yes | 0 | 0 | 1 | $2.40 \cdot 10^{-6}$ |
| Pfu | Yes | 0 | 0 | 1 | $2.80 \cdot 10^{-6}$ |
| **Storage Host** | **Domain** |  |  |  |  |
| *E. coli* | Prokaryotes | 0.08 | 0.08 | 0.84 | $3.17 \cdot 10^{-7}$ |
| *H. sapiens* | Eukaryotes | 0.06 | 0.06 | 0.88 | $6.90 \cdot 10^{-11}$ |
| *M. musculus* | Eukaryotes | 0.025 | 0.025 | 0.95 | $4.40 \cdot 10^{-9}$ |
| *D. melanogaster* | Eukaryotes | 0.33 | 0.33 | 0.34 | $2.10 \cdot 10^{-8}$ |
| *S. cerevisiae* | Eukaryotes | 0.13 | 0.13 | 0.74 | $7.90 \cdot 10^{-8}$ |
| **Sequencing method** | **Submethod** |  |  |  |  |
| Illumina | Single end | 0.0024 | 0.0013 | 0.9963 | $2.10 \cdot 10^{-3}$ |
| Illumina | Paired end | 0.0018 | 0.0011 | 0.79 | $3.20 \cdot 10^{-3}$ |
| Nanopore | 1D | 0.37 | 0.15 | 0.48 | $2.00 \cdot 10^{-1}$ |
| Nanopore | 2D | 0.36 | 0.23 | 0.41 | $1.30 \cdot 10^{-1}$ |
| PacBio | Subread | 0.2 | 0.05 | 0.75 | $2.00 \cdot 10^{-1}$ |
| PacBio | CCS | 0.21 | 0.42 | 0.37 | $1.40 \cdot 10^{-1}$ |

### 2.2 Framework structure

All methods, optional components, and Autoturbo-DNA hyperparameters are embedded into a PyTorch framework. Each option can be selected and changed using command line arguments or by supplying a configuration file. Error sources and their properties, as well as error probabilities for breaches of constraints, can be supplied in JSON files. After each epoch, the current optimizer state, model, configuration file, and a log file containing the loss value of each component, as well as the accuracy, stability, noise level, and percentage of correctly recovered blocks, are saved. The stability is defined as

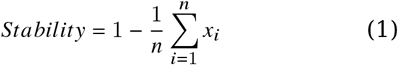

with *x*_*i*_ being the error probability of the *i*th coded block of batch size *n*. The error probability is the user-defined probability of an error occurring in the storage channel due to constraint breaches, i.e., GC content that is not in the range set by the user or homopolymer chains longer than desired. The training can be paused anytime and will resume after the last evaluated epoch. Hyperparameters that are not integral to the model structure, like the model type, number of layers, or activation functions, can be adjusted during training, allowing online hyperparameter tuning.

### 2.3 Autoencoder structure

#### Interleaver

The main codec combines three main components, as shown in Figure 1. Multiple options are available for each component, as shown in supplemental tables 2 to 4. One of the fundamental parts of turbo codes, and, by extension, turbo autoencoders, is the interleaver. The interleaver permutes the input sequence using either a pseudo-random function, with the seed of the function known to both the encoder and the decoder, or a deterministic interleaver with the geneneral form [16] of

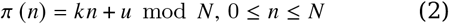

with *k* and *n* as fixed integers, and *k* being relatively prime to *n*. In a traditional turbo code design, the interleaver serves to spread out burst errors, improving the convergence of the decoding algorithm [16]. For turbo autoencoders, which are primarily evaluated using i.i.d channels, the interleaver adds long-range memory to the code structure instead of increasing its robustness against burst errors [13]. For Autoturbo-DNA, both the addition of long-range memory and the increased robustness of the code against burst errors by the application of an interleaver is of importance, as indel errors, which affect all bases following the error, can be interpreted as burst errors.

**Figure 1:**
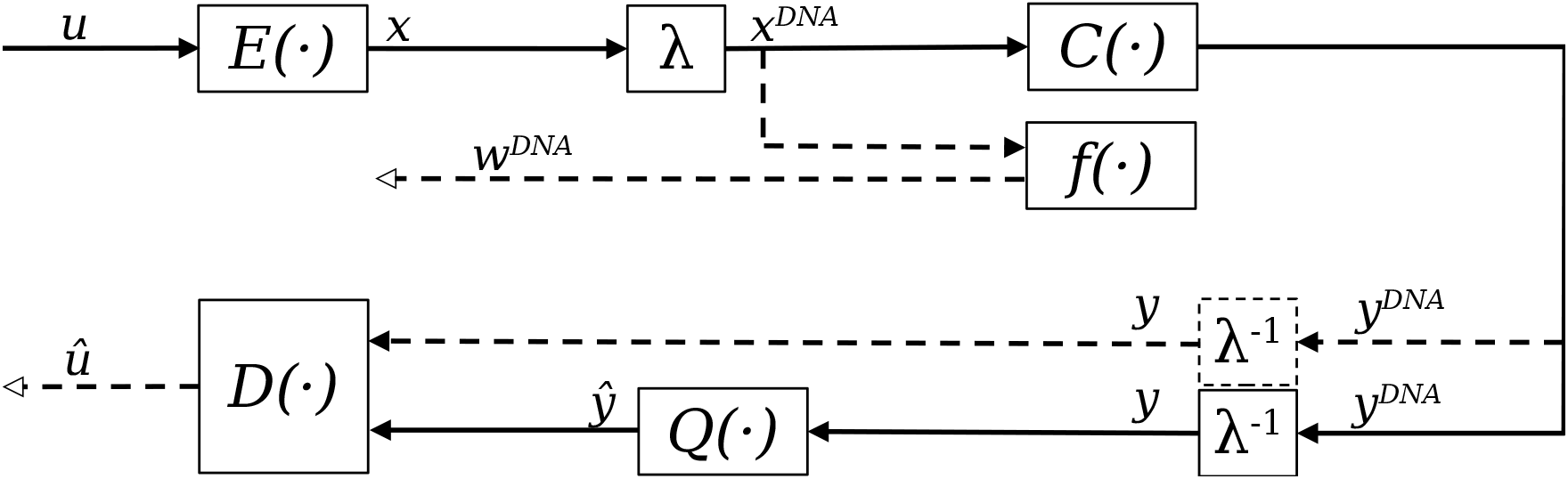
Overview of the main components of Autoturbo-DNA: A given binary input *u* is encoded using the encoder network *E* and subsequently mapped to a DNA sequence by the mapping function *λ*. The output of the mapping function *x*^*DN A*^ is then modified by the channel simulator *C*(·) and optional evaluated for constraint adherence by the function *f*(·) . The output score *w*^*DN A*^ of the evaluation function can be used as an additional loss metric for the encoder. The inverse of the mapping function translates the DNA sequence back into a binary representation *y*. Depending on the training stage and chosen configuration, the channel output *y*^*DN A*^ is either directly decoded by the decoder function *D*(·) or first transcoded using the indel reduction component *Q*(·), and the resulting sequence *ŷ* is decoded by the decoder function, producing the binary output sequence *û*.

#### Encoder

Each of the supported encoder network structures shown in supplementary table 2 supports a base code rate of either 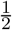 by encoding one copy of the input data without interleaving and one copy that is interleaved before being encoded, or 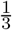, in which two copies of the input data are first encoded separately without interleaving. One copy is interleaved before the encoding. The encoders further support more granular adjustments to the code rate by increasing neurons in the output layer. The basic structure of the encoder is shown in Figure 2. The output is further normalized as described in [17, 13]:

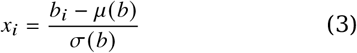

with *μ* (*b*) and *σ* (*b*), respectively, being the mean and standard deviation of the block. As the output of the encoder has to be mapped to the four DNA nucleotides, *x*^*DN A*^ ⊆{*A, T, C, G*}, the output of the encoder is binarized and combined with a straight-through estimator as described in [13]. Each separately encoded copy of the input data is, after encoding, concatenated with each other. A mapping function *λ* maps each bit pair to a base, with 0 ∧ 0 = *A*, 0 ∧ 1 = *G*, 1 ∧ 0 = *T*, and 1 ∧ 1 = *C*. The encoder is trained, either separately or in conjunction with the other parts of the codec, using the smooth L1 loss between the input data and the output of the decoder:

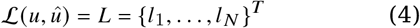

**Figure 2:**
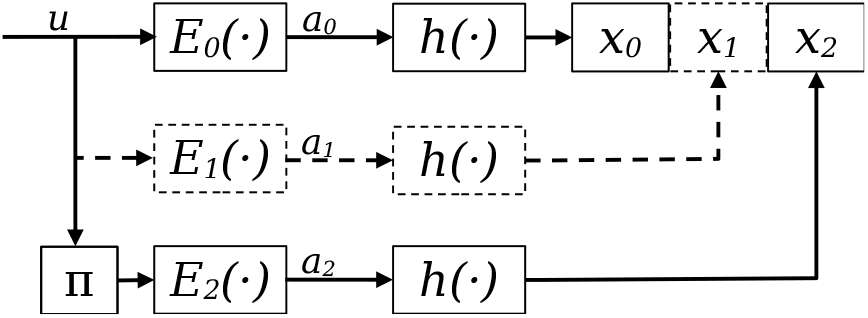
Schematic illustration of the encoder: It consists of two, with an optional third, networks *E*_0− 2_(·) . The input data is copied and sent through each encoder network separately. For the encoder *E*_2_(·), the input data is first interleaved using the interleaving function *π* before being encoded. Each encoder output is normalized and binarized before being concatenated into the encoded binary sequence *x*.

For a batch size of *N*, with

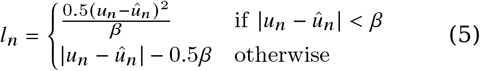

The hyperparameter *β* is user-definable, with a default value 1.0.

### Constraint adherence training

Additionally to training the encoder for error correction performance, it can also be trained to encode the data into a constraint-adhering representation. Autoturbo-DNA supports the constraints of GC content, homopolymer length, k-mer occurrence, and undesired motifs. The undesired motifs are supplied using a JSON file that contains for each undesired motif an entry consisting of the motif itself, the error probability that is associated with the occurrence of the motif, and, optionally, a description of the motif. The GC content, homopolymer length, and k-mer occurrence are defined in a JSON file as a graph’s x and y coordinates. For example, a GC content that has a 100 % error probability between 0 % and 40 % GC content and between 60 % and 100 %, with the area between 40 % and 60 % having a 0 % error probability, can be described with the points X = 0, Y = 100; X = 40, Y = 100; X = 41, Y = 0; X = 59, Y = 0; X = 60, Y = 100, and X = 100, Y = 100. This approach further allows interpolation to smooth curves between points. Besides manual editing, the JSON file, error probability plots for the GC content, homopolymer length, and k-mer occurrence can be created using the graphical interface of MESA, as Autoturbo-DNA is cross-compatible with the configuration files of MESA. This cross-compatibility allows the generation of complex patterns by dragging points of a graph or simply combining common undesired motifs from a large selection of pre-existing ones in MESA.

### Indel reduction component

Upstream of the decoder *D* (·) is the indel reduction component *Q* (·) . As the decoder requires a fixed-size input, the primary purpose of this component is to transform the noisy sequences into a fixed-size representation that can be used as input for the decoder. It further serves as a transcoder that maps the quaternary DNA sequences to either binary sequences or sequences of continuous values. Autoturbo-DNA supports a multitude of strategies for indel reduction, as shown in supplementary table 3. By default, it is trained using the smooth L1 loss of the encoder output with the output of *Q* (·) . The output of *Q* (·) is binarized to allow for this training. Alternatively, it is also possible to train the indel reduction component using the smooth L1 loss between the input and decoded data. This approach does not require binarization and the continuous output of *Q* (·) can be passed to the decoder.

### Decoder

The decoder component follows the principles established by.^13^ It comprises two decoders, concatenated in serial, that iteratively update their prediction by utilizing the posterior of the previous decoder as prior.

For a given input sequence, the sequence is split into multiple sub-sequences corresponding to the encoder’s two or three output sequences. The amount of sequences the input is split into depends on the chosen rate, with two sequences for a rate of 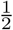 and three sequences for a rate of 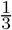 . For a rate of 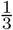, *y*_0_ and *y*_1_ are used as input to the first decoder, with a prior initialized as a tensor consisting only of zeros, the same size as the input subsequences. The output of the first decoder is then transposed, using the interleaving function with the same seed as used by the encoder, leading to the same inter-leaving pattern. Besides the posterior of the first decoder, the second decoder also takes as input *y*_2_, corresponding to the interleaved encoded subsequence, and *y*_0_, interleaved using the same seed as the encoder. The inverse function of the inter-leaver is then applied to the posterior of the second decoder, which is subsequently passed to the first decoder as prior. This sequence is repeated until a user-definable amount of iterations is reached. The output of the final iteration is then passed to a sigmoid activation function, resulting in the final output sequence. A schematic of the decoding process for a rate of 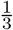 is shown in Figure 3 for a rate of 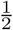. The input of the first decoder is *y*0 and the deinterleaved *y*_1_, together with the prior, as described above. In contrast, the input for the second decoder is the interleaved form of *y*_0_, together with *y*_1_ and the interleaved prior.

**Figure 3:**
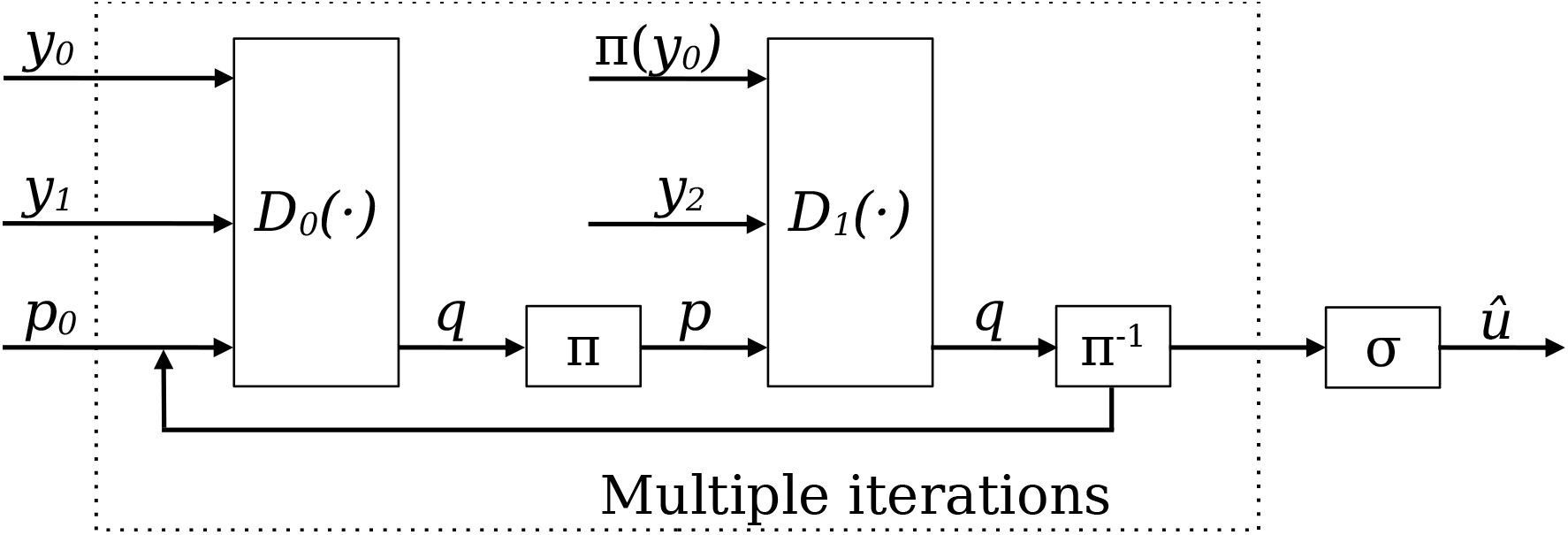
Schematic illustration of the decoder, shown for rate 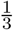. The input sequence *y* is split into the subsequences *y*_0_, *y*_1_ and *y*_2_, which correspond to the encoded subsequences *x*_0_, *x*_1_ and *x*_2_, respectively. *y*_0_ and *y*_1_, together with a prior *p*_0_, are used as the input for the first decoder network *D*_0_ (·). The output posterior *q* and *y*_0_ are subsequently interleaved and serve, together with *y*_2_, as the input for the second decoder *D*_1_ (·) . The output of the second decoder is deinterleaved by the inverse of the interleaving function, *π* ^(^ −1), and is used as updated prior for the first decoder. This process will be repeated until a user-defined amount of iterations is reached. The final, deinterleaved output of the second decoder will then be used to generate the output sequence *û* by a sigmoid activation function *σ*.

## 3 Results

To showcase the versatility and flexibility of Autoturbo-DNA, we have trained models with different combinations of encoder architectures, transcoder architectures, and decoder architectures. This approach aims to isolate the effects of different encoder-decoder combinations on the overall performance and outcomes, providing an indicative illustration of the potential usage of the framework. To achieve this, we have set a fixed set of hyperparameters across all trials to give a snapshot of the framework’s functionality under constant conditions. The hyperparameters that deviate from the default values are shown in table 2. The error probabilities used are a combination of column-synthesized oligos utilizing the ErrASE error correction, Illumina single-end sequencing, PCR amplification for 30 cycles using the *T AQ* polymerase, and storage in *E* .*coli* for 24 months. This combination of error sources would lead to a total error rate of 0.33 %. The error rate was increased while keeping the distribution of error types and patterns, using the amplifier to a total error rate of 6.8 %, which is closer to the error rate of other DNA data storage codes used for in-vitro storage and which include constraint adherence [10].

**Table 2:** Hyperparameters used in the evaluations that deviate from the default Autoturbo-DNA hyperparameters.

| Parameter | Value |
| --- | --- |
| Amplifier | 15 |
| Epochs | 400 |
| Encoder dropout | 0.2 |
| Encoder layers | 7 |
| Encoder learning rate | 0.00001 |
| Encoder steps | 20 |
| Decoder layers | 7 |
| Decoder learning rate | 0.00001 |
| Decoder steps | 30 |
| Decoder iterations | 10 |
| Transcoder layers | 7 |
| Transcoder learning rate | 0.00001 |
| Transcoder steps | 20 |
| Transcoder-units | 128 |
| Transcoder train target | Encoded data |
| Combined steps | 20 |
| Weight initialization | Normal |
| Block length | 8 |

### 3.1 Architecture comparison

To compare the different encoder, transcoder, and decoder architectures, the reconstruction accuracy of each model is calculated and shown as box-plots. For the different encoders, the results are shown in Figure 4. The results indicate that models utilizing either CNNs, VAEs, or RNNs as encoder architecture perform similarly well for the used hyperparameters and evaluation metrics. At the same time, a transformer-encoder has the lowest mean and median from all evaluated encoder architectures.

**Figure 4:**
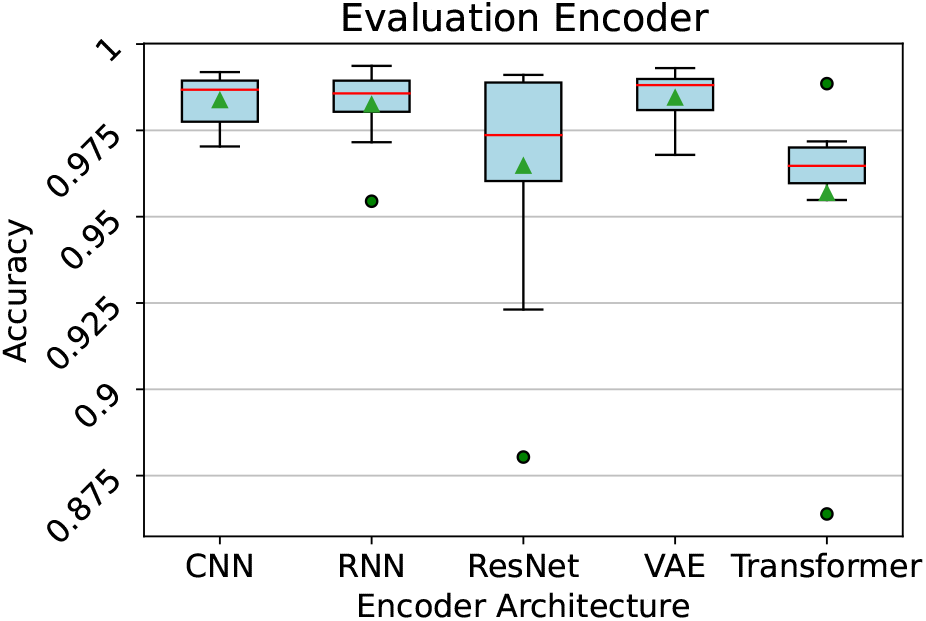
Boxplot of the accuracy of the trained models, separated by the used encoder architecture. A red line represents the median, a green triangle represents the mean, and the outliers are represented by green dots.

ResNets as transcoder architectures have the highest mean and median and the smallest interquartile range for the evaluated hyperparameters and evaluation metrics, as shown in Figure 5. RNNs as transcoder architecture led to a median accuracy that is slightly higher than the median accuracy of CNNs. However, RNNs have a lower mean and a more extensive interquartile range than CNNs and RNNs.

**Figure 5:**
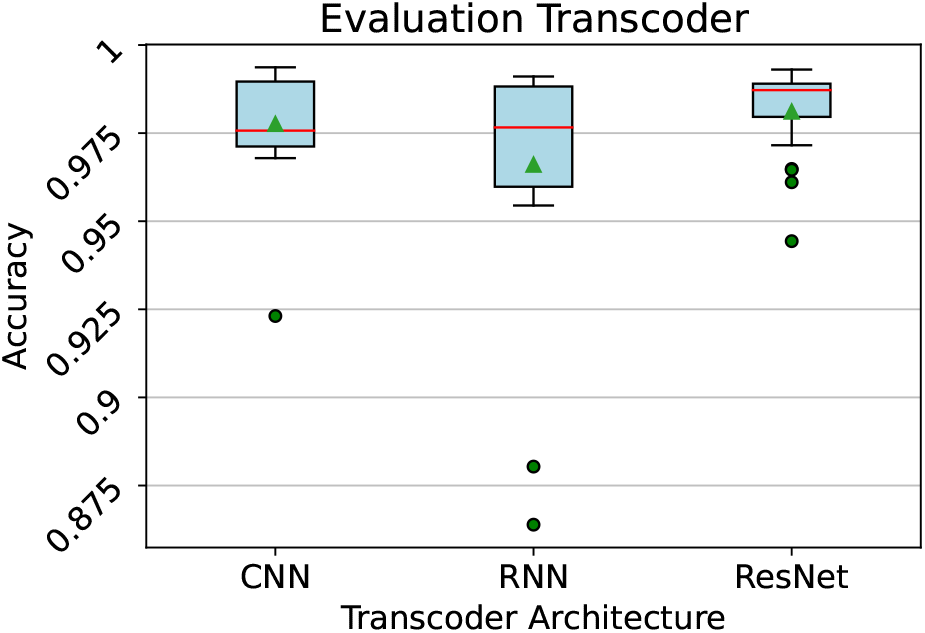
Boxplot of the accuracy of the trained models, separated by the used transcoder architecture. A red line represents the median, a green triangle represents the mean, and the outlier are represented by green dots.

For the decoder architectures, shown in Figure 6, CNNs had the highest mean and median accuracy and the smallest interquartile range. ResNets had a median accuracy close to CNNs but with a larger interquartile range, lower mean accuracy, and less outliers. Transformer-based encoders and RNNs as decoder architectures performed similarly with the chosen evaluation metrics and hyperparameters.

**Figure 6:**
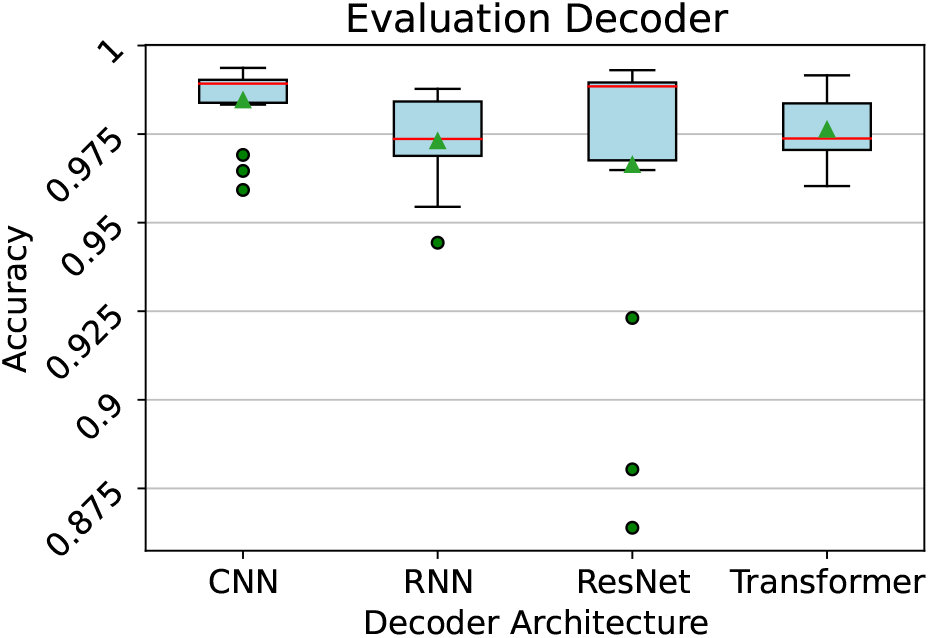
Boxplot of the accuracy of the trained models, separated by the used decoder architecture. A red line represents the median, a green triangle represents the mean, and the outlier are represented by green dots.

For further evaluations, we focused on VAE and CNN architectures for the encoder, CNN architectures for the decoder, and ResNets for the transcoder.

### 3.2 Block length comparison

To evaluate the influence of the block length on the training time and reconstruction accuracy, we used the same set of parameters described in Section 3.1. However, we increased the number of epochs to 1,000 to ensure the models have sufficient time to learn and extract features from the more complex, longer sequences, thereby effectively capturing their underlying patterns and structures. We used block lengths of 3 ·8, 3 ·16, 3 · 32, and 3· 64 bits. The results of the reconstruction accuracy over time, with a rolling average over ten epochs, are shown in Figure 7. The results without the rolling average and results that compare the encoder architectures with different block lengths are shown in the Supplement.

**Figure 7:**
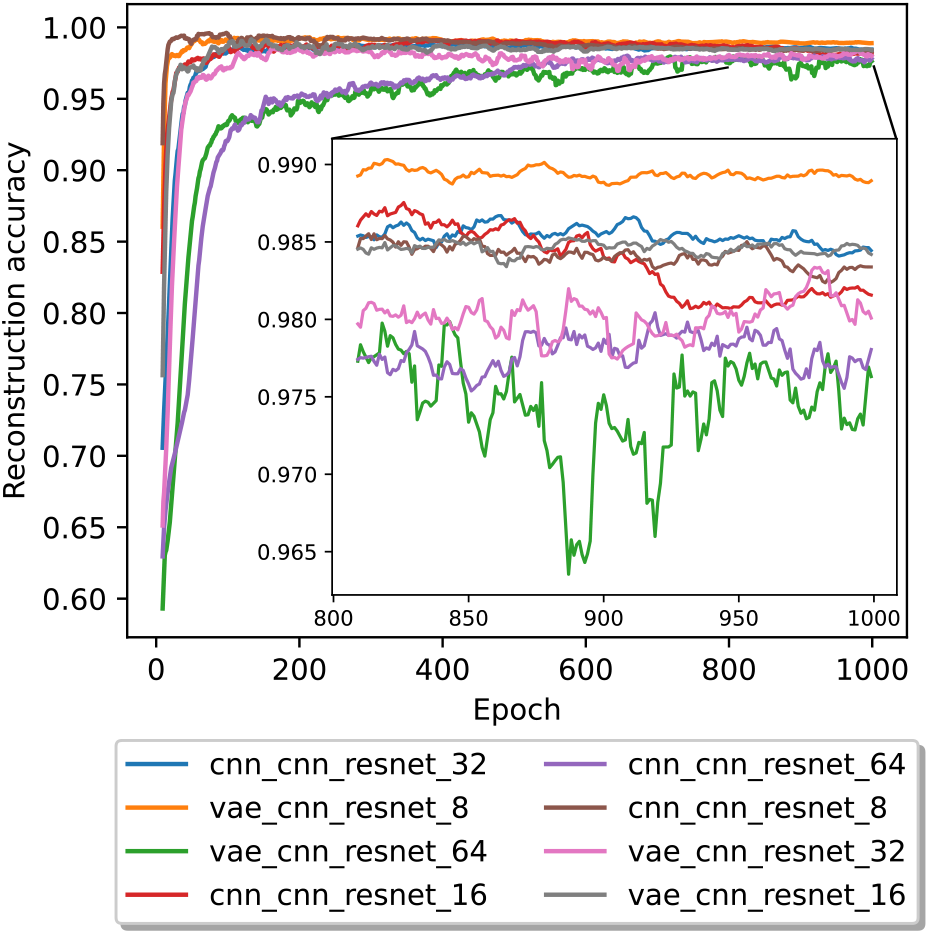
The average reconstruction accuracy in a 10 epoch rolling window for 1,000 epochs. For visual clarity, the last 200 epochs are magnified. The legend labels are structured in the form of encoder, decoder, transcoder, block size.

With shorter block lengths, the reconstruction accuracy increased in fewer epochs than with larger block lengths and also reached higher peaks. With a block length of 3·8 bits, a model using a CNN encoder reached a peak reconstruction accuracy of 0.99744 after 90 epochs. In comparison, the VAE encoder model reached a peak of 0.99576 after 170 epochs. The CNN encoder-based model showed a clear downtrend in reconstruction accuracy over time (as shown in the supplement). With a block length of 3 · 16 bits, a CNN encoder model reached a peak reconstruction accuracy of 0.99318 after 438 epochs, and the VAE encoder model reached a peak reconstruction accuracy of 0.99455 after 148 epochs. With this block length, CNN encoder-based models also showed a down-trend in reconstruction accuracy over time. For a block length of 3 · 32 bits, the CNN encoder model peaked at a reconstruction accuracy of 0.99021 after 105 epochs, followed by a more stable progression than the shorter block lengths. The VAE encoder model reached a peak of 0.9901 reconstruction accuracy after 161 epochs. In contrast, with a block length of 3 · 64 bits, the CNN encoder model reached its peak of 0.984 reconstruction accuracy after 604 epochs, and the VAE encoder model reached its peak of 0.98452 after 937 epochs.

### 3.3 Impact of latent redundancy

Besides supporting the rates 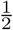 and 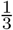 by changing the number of encoded blocks that are generated per block of input data, Autoturbo-DNA also supports more fine-grading adjustments of the code rate, by increasing the number of units of the encoder layers, leading to an increase in the size of the latent representation. To analyze the impact of an increase in latent representation size, the same hyperparameters as before were used, but with the latent redundancy hyperparameter set to either 2 bits, 4 bits, or 8 bits and with a block length of either 8 or 16 bits. The results of the average reconstruction accuracy, from epoch 200 on, and for a block length of 8 bits are shown in Figure 8. The results for all 1,000 epochs, as well as results without the rolling average, are shown in the supplement.

**Figure 8:**
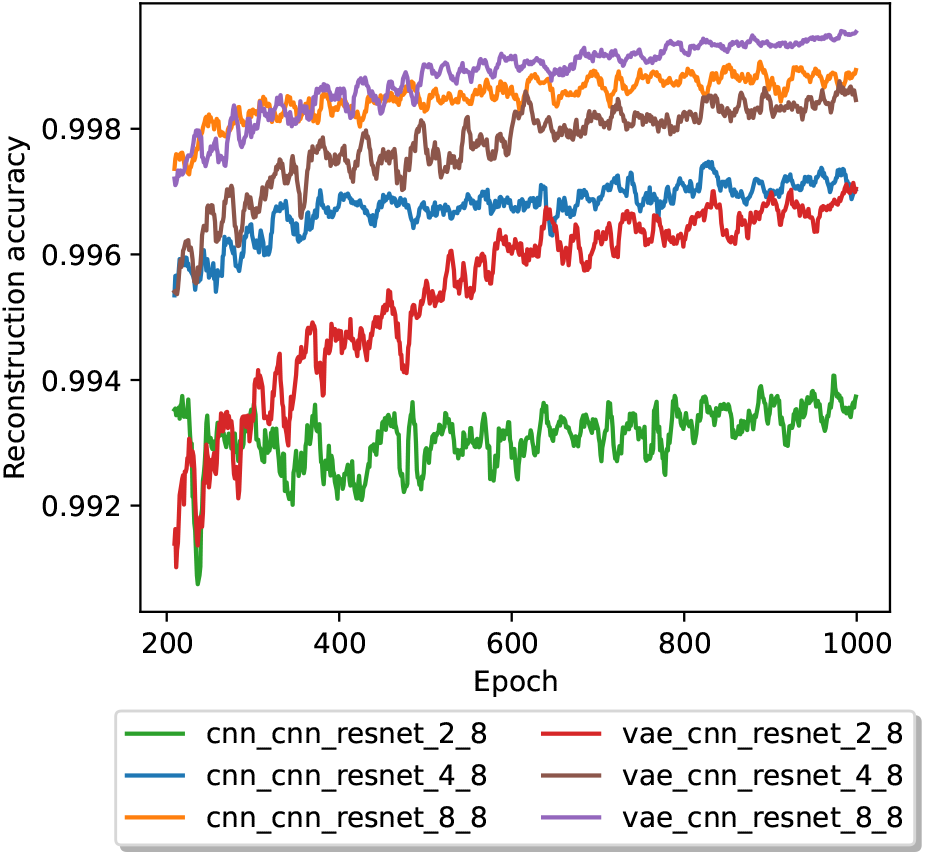
The average reconstruction accuracy in a 10 epoch rolling window for 1,000 epochs, beginning with epoch 200 and with a block size of 3·8. The legend labels are structured in the form of encoder, decoder, transcoder, latent redundancy, block size.

In contrast to the results without latent redundancy, no model showed signs of a downtrend. Each model had a higher reconstruction accuracy than any model without latent redundancy. For the CNN model with a latent redundancy of 2 bits, the reconstruction accuracy was 0.99442, while for the VAE model with the same amount of redundancy, the accuracy was 0.99725. For a latent redundancy of 4 bits, the CNN model had a accuracy of 0.99683, while the VAE model had a accuracy of 0.99817. For an 8-bit latent redundancy, the CNN model had a accuracy of 0.99899, and the VAE model had a accuracy of 0.99957. For each latent redundancy value, the accuracies of the models utilizing a VAE-based encoder are higher than those based on a CNN encoder. For all tested models and latent redundancy values, the slopes of the last 200 epochs are slightly positive, as shown in Table 3.

**Table 3:**
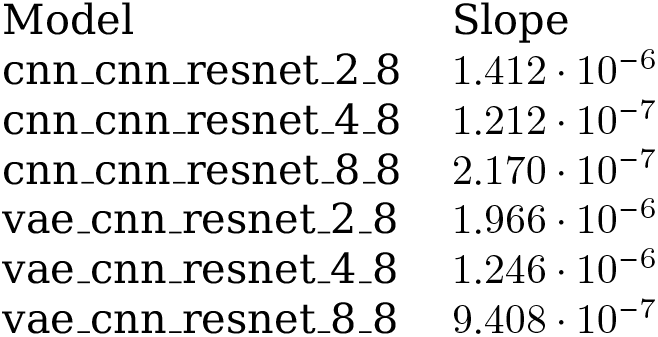
Slopes of the last 200 epochs for the evaluated models using a block size of 3 · 8. The naming convention is encoder, decoder, transcoder, latent redundancy, block length.

**Table 4:** Slopes of the last 200 epochs for the evaluated models using a block size of 3 · 16. The naming convention is encoder, decoder, transcoder, latent redundancy, block length.

| Model | Slope |
| --- | --- |
| cnn_cnn_resnet_2_16 | $1.956 \cdot 10^{-6}$ |
| cnn_cnn_resnet_4_16 | $-2.422 \cdot 10^{-7}$ |
| cnn_cnn_resnet_8_16 | $4.868 \cdot 10^{-6}$ |
| vae_cnn_resnet_2_16 | $2.878 \cdot 10^{-6}$ |
| vae_cnn_resnet_4_16 | $2.377 \cdot 10^{-6}$ |
| vae_cnn_resnet_8_16 | $2.944 \cdot 10^{-6}$ |

Using a block length of 3 · 16, the reconstruction accuracy was lower for each encoder architecture and latent redundancy combination than the same combinations with a block size of 3· 8. For the models using a CNN encoder, the accuracies were as follows: 0.99025 for a latent redundancy of 2 bits, 0.99272 accuracy for a latent redundancy of 4 bits, and an accuracy of 0.99545 for a latent redundancy of 8 bits. Utilizing a VAE-based encoder, the accuracies were 0.98605 for a latent redundancy of 2 bits, 0.98886 for a latent redundancy of 4 bits, and 0.99377 for a latent redundancy of 8 bits. In contrast to the evaluations where a block size of 3 · 8 was used, with a block size of 3 · 16, a CNN-based encoder leads to a higher accuracy score for all tested latent redundancies. Except for the CNN-based encoder utilizing a latent redundancy of 4 bits, all models had a slight positive slope for the last 200 epochs.

### 3.4 Fine-tuning for constraint adherence

To train the encoder to generate DNA sequences that adhere to constraints, we have generated a configuration file using MESA that associates a 100 % error probability to a sequence if it contains a homopolymer longer than three bases or if the GC content of the sequence is not between 40 % to 60 %. These constraints were chosen for comparability to other studies 10. The stability score, which is used as an additional metric to train the encoder in addition to the reconstruction accuracy, is calculated from the sum of the error probability *e* of each sequence for a batch size of *N* in the following way:

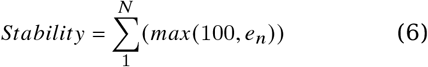

Training the models from the beginning using the stability score as additional training metric for the encoder has only a slight impact on the stability of the stability score of the models, as shown in Figure 10, with the impact on the complete training period and on the accuracy shown in the supplement.

Instead of training the models from the beginning using the additional constraint adherence training, we also fine-tuned the models that were trained with different amounts of latent redundancy (see Figure 9 and Figure 10). The fine-tuning was carried out and led to a significant (t-statistic: 2.91, p-value: 0.008) difference in the means between the two groups, as shown in Figure 11 and over the complete course of the training in the supplement. After fine-tuning, the mean and median of the accuracies were slightly lower than before, as shown in the supplement. The difference in means was, however, not statistically significant, with a t-statistic of -0.932 and a p-value of 0.362.

**Figure 9:**
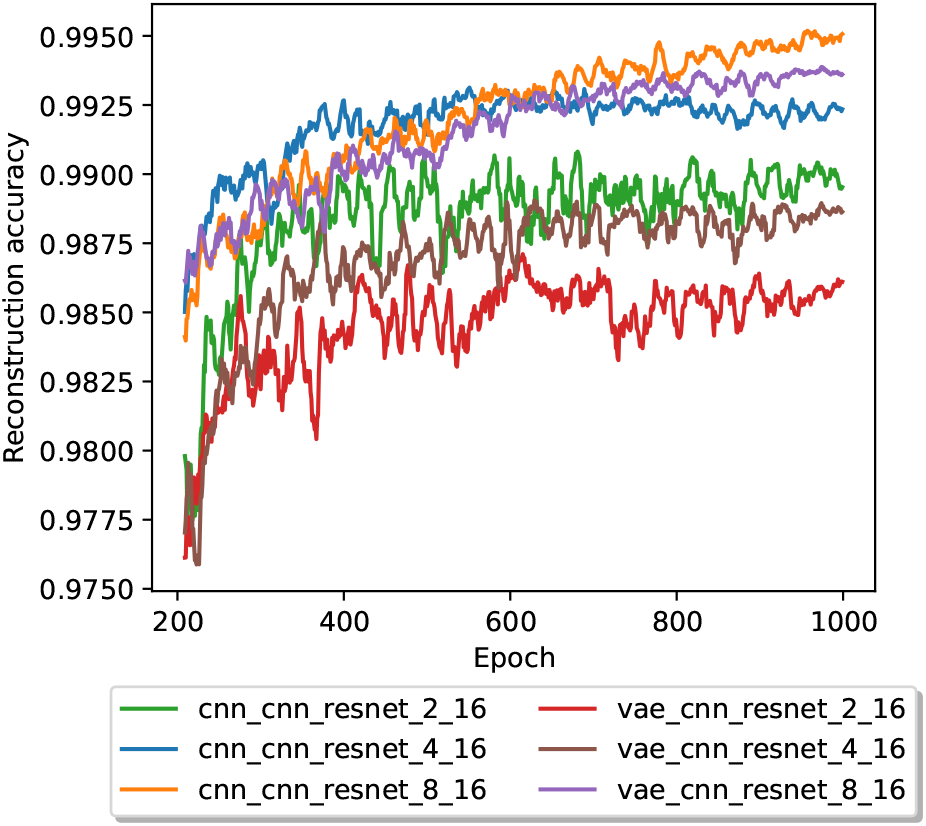
The average reconstruction accuracy in a 10 epoch rolling window for 1000 epochs, beginning with epoch 200 and with a block size of 3 · 16. The legend labels are structured in the form of encoder, decoder, transcoder, latent redundancy, block size.

**Figure 10:**
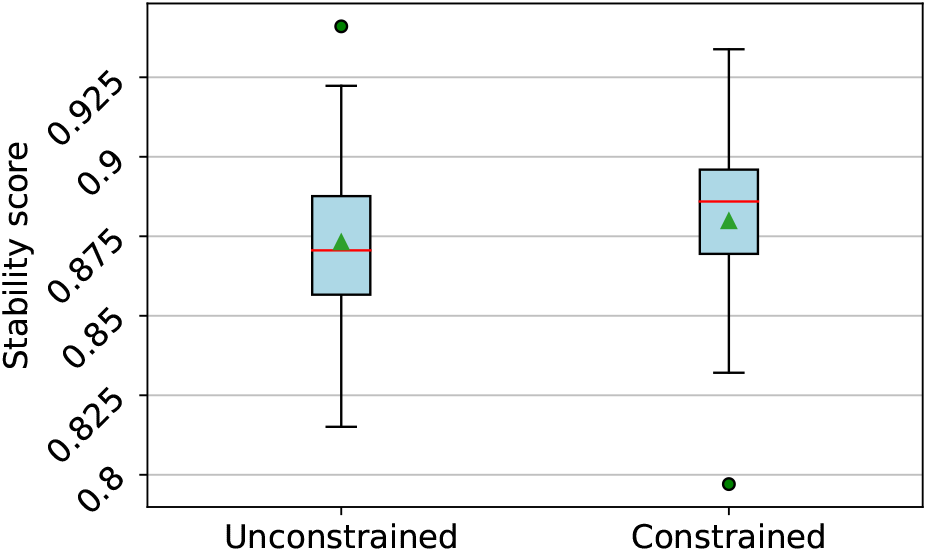
Boxplot of the stability score of models trained without (left) and with (right) the stability score as a training metric. The models were further trained with either 2, 4, or 8 bits of latent redundancy and a block size of 8 or 16 bits. A red line represents the median, a green triangle represents the mean, and the outlier are represented by green dots.

**Figure 11:**
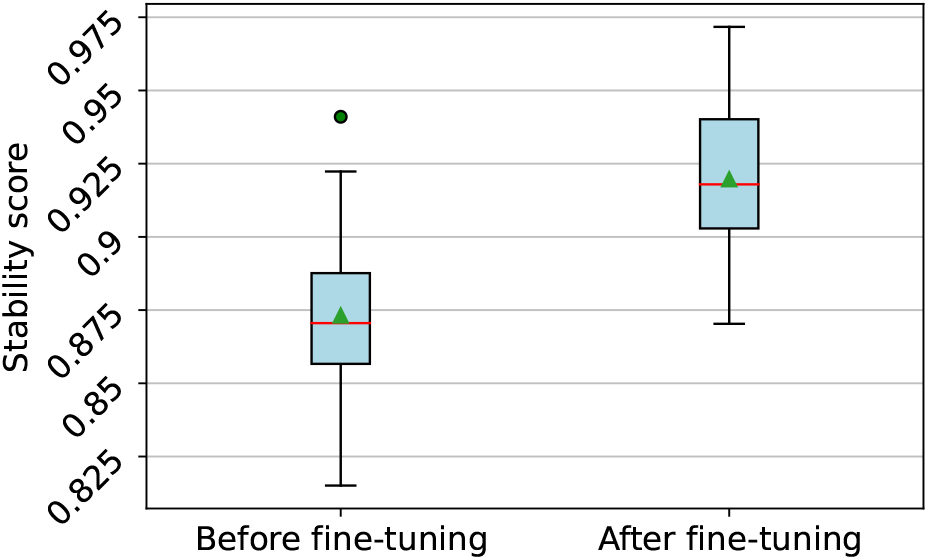
Boxplot of the stability score of models trained before (left) and after (right) fine-tuning for 100 epochs using the stability score as a training metric. The models were further trained with either 2, 4, or 8 bits of latent redundancy and a block size of 8 or 16 bits. A red line represents the median, a green triangle represents the mean, and the outlier are represented by green dots.

## 4 Discussion

Autoturbo-DNA is a feature-rich framework for training encoder-transcoder-decoder models for DNA data storage. The framework supports training using a DNA data storage channel simulator with a wide variety of options, all based on literature error-rates and -patterns of the individual DNA storage channel components. Utilizing the TurboAE structure first presented by [13], combined with a transcoder to account for insertion and deletion errors, allows Autoturbo-DNA to reach reconstruction performance close to single sequence non-neural-network state-of-the-art error correction and constrained codes for DNA data storage [10]. The results for the evaluations including latent redundancy bits indicate potential further improvements with longer training time, while a high number of constraint free sequences can be generated by fine-tuning the model by using the stability score as an additional metric to train the encoder on.

Further potential improvements in error correction and constraint adherence performance could be achieved by optimizing the hyperparameters for different encoder, transcoder, and decoder combinations. Additional, multiple potential improvements to the basic TurboAE implementation are described, which could potentially be used to improve the performance of Autoturbo-DNA, for example, utilizing a trainable interleaver [18, 19], or more sophisticated training strategies [20]. Our results indicate that neural-network based codecs could be a viable alternative to traditional codecs for the DNA data storage channel.

## Code availability

The Autoturbo-DNA framework is available at https://github.com/MW55/autoturbo_dna. The framework is available under the MIT license.

## Acknowledgements

The authors want to thank Hannah Franziska Löchel for fruitful discussions regarding the figure design. This work was financially supported by the LOEWE program of the State of Hesse (Germany) in the MOSLA research cluster. This work was supported by the BMBF-funded de.NBI Cloud within the German Network for Bioinformatics Infrastructure (de.NBI) (031A532B, 031A533A, 031A533B, 031A534A, 031A535A, 031A537A, 031A537B, 031A537C, 031A537D, 031A538A).

## Author Contributions Statement

MW conceived the initial idea. HD developed the initial version of the framework under the supervision of MW. MW developed the final version of the framework. MW wrote the initial draft, with contributions from DH. MW created the figures, with contributions from HD. MW carried out the evaluations. DH supervised the study. All authors contributed to the final manuscript.

