## Supplemental information for "Autoturbo-DNA: Turbo-Autoencoders for the DNA data storage channel"

September 2023

### 1 Configuration parameters

| Option Strings | Type | Default | Help |
|---|---|---|---|
| -h, –help | None | ==SUPPRESS== | show this help message and exit |
| -v, –version | None | ==SUPPRESS== | show program's version number and exit |
| –wdir | str | None | Path to the working directory, if not existing the model will be saved here, if already existing the model will be loaded. |
| –train | bool | False | Create and train the desired model. |
| –bitenc | None | None | Encode with a model a bit string into a code. |
| –bitdec | None | None | Decode with a model a bit string into a code. |
| –encode, -e | None | None | Encode with a model a file. |
| –decode, -d | None | None | Decode with a model a code back into a file. |
| –input, -i | str | None | Path to the file to be en-/decoded. |
| –output, -o | str | None | Path to the output file. |
| –index_size, -is | int | 16 | size (in bits) of the added index, larger files need bigger index sizes, has to be a multiple of 8. |
| –simulate | None | None | Simulate errors on a generated code. |
| –ids | None | False | Shows a list of the default ids of the different options for DNA synthesis, storage and sequencing simulation. |
| –seed | int | 0 | Specify a integer number, this allows to reproduce the results. |
| –gpu | None | False | Whether the calculations of the models should run on the GPU (using CUDA). |
| –parallel | None | False | Whether to run the calculations on multiple GPUs, if there are more than one. |
| –threads | int | 8 | If using the CPU, how many threads should be used. |
| –rate | str | onethird | Rate of the code, supported are 1/3 (argument=onethird) and 1/2 (argument=onehalf) |
| –block-length | int | 64 | Length of the bitstreams to be used |
| –block-padding | int | 18 | Length of the padding by which the bitstream is extended |

Continued on next page

| Option Strings | Type | Default | Help |
|---|---|---|---|
| –encoder | str | cnn | Choose which encoder to use: RNN, SRNN, CNN, SCNN or RNNatt |
| –enc-units | int | 64 | The number of expected features in the hidden layer for the encoder |
| –enc-actf | str | elu | Choose which activation function should be applied to the encoder: tanh, elu, relu, selu, sigmoid or identity |
| –enc-dropout | float | 0.0 | Dropout probability for the encoder |
| –enc-layers | int | 5 | Number of recurrent layers per RNN/CNN structure in the encoder |
| –enc-kernel | int | 5 | Size of the kernels for the CNN in the encoder |
| –enc-rnn | str | GRU | Choose which structure to use for the RNN in the encoder: GRU or LSTM |
| –vae-beta | float | 0.0 | The beta multiplier of the Kullback–Leibler divergence if using a VAE. |
| –decoder | str | cnn | Choose which decoder to use: RNN or CNN |
| –dec-units | int | 64 | The number of expected features in the hidden layer for the decoder |
| –dec-actf | str | identity | Choose which activation function should be applied to the decoder: tanh, elu, relu, selu, sigmoid or identity |
| –dec-dropout | float | 0.0 | Dropout probability for the decoder |
| –dec-layers | int | 5 | Number of recurrent layers per RNN/CNN structure in the decoder |
| –dec-inputs | int | 5 | The number of expected input features for the decoder |
| –dec-iterations | int | 6 | Number of iterative loops to be made in the decoder |
| –dec-kernel | int | 5 | Size of the kernels for the CNN in the decoder |
| –dec-rnn | str | GRU | Choose which structure to use for the RNN in the decoder: GRU or LSTM |
| –not-extrinsic | None | True | Whether extrinsic information should be applied to the decoder each iteration |
| –coder | str | cnn | Choose which coder to use: MLP, CNN or RNN |
| –coder-units | int | 64 | The number of expected features in the hidden layer for the coder |
| –coder-actf | str | elu | Choose which activation function should be applied to the coder: tanh, elu, relu, selu, sigmoid or identity |
| –coder-dropout | float | 0.0 | Dropout probability for the coder |
| –coder-layers | int | 5 | Number of recurrent layers per RNN/CNN structure in the coder |
| –coder-kernel | int | 5 | Size of the kernels for the CNN in the coder |
| –coder-rnn | str | GRU | Choose which structure to use for the RNN in the coder: GRU or LSTM |
| –init-weights | str | None | Choose which method to use to initialize the linear layers of the model: normal, uniform, constant, xavier_normal, xavier_uniform, kaiming_normal or kaiming_uniform |

Continued on next page

| Option Strings | Type | Default | Help |
|---|---|---|---|
| –lat-redundancy | int | 0 | Redundancy of the final encoder layer (and first decoder layer), required to account for constraints. Has to be divisible by 2 |
| –ens-models | int | 3 | If ensemble coders are used, defines the number of coder instances in the ensemble. |
| –padding-style | str | constant | If padding should be constant values or a circular copy of the input. |
| –blocks | int | 1024 | Number of the bitstreams to be used |
| –batch-size | int | 256 | Size of the batch to be used during training |
| –epochs | int | 100 | Number of epochs the whole model should be trained |
| –enc-lr | float | 0.00001 | Value of the learning rate to be used for the encoder |
| –enc-optimizer | str | adam | Choose which optimizer to use for the encoder: Adam, SGD or Adagrad |
| –enc-steps | int | 1 | Number of training steps to be performed per epoch for the encoder |
| –dec-lr | float | 0.00001 | Value of the learning rate to be used for the decoder |
| –dec-optimizer | str | adam | Choose which optimizer to use for the decoder: Adam, SGD or Adagrad |
| –dec-steps | int | 2 | Number of training steps to be performed per epoch for the decoder |
| –coder-lr | float | 0.001 | Value of the learning rate to be used for the coder |
| –coder-optimizer | str | adam | Choose which optimizer to use for the coder: Adam, SGD or Adagrad |
| –coder-steps | int | 5 | Number of training steps to be performed per epoch for the coder |
| –simultaneously | None | False | Whether the encoder and decoder are to be trained at the same time, if so, the learning parameters from the encoder are used |
| –batch-norm | bool | False | Whether to use batch normalization or not. |
| –separate-coder-training | None | False | If the coder should be split into 3 seperate instances during training. |
| –all-errors | None | False | train each part of the model always with all error types. |
| –channel | str | dna | which channel model should be used for training |
| –continuous-coder | None | False | toggles that the intermediate decoder (coder) passes continuous values to the decoder. |
| –constraint-training | None | False | If the code should also be trained to adhere to constraints. |
| –loss-beta | float | 1.0 | beta parameter for the smooth L1 loss. |
| –coder-train-target | str | encoded_data | how the coder should be trained, for best reconstruction accuracy or to be as close to the encoder output as possible |
| –simultaneously-warmup | int | 0 | if using simultaneously training, how many warmup epochs should be trained seperatly, before moving to simultaneously training. |
| –synthesis | None | (1, None) | Specify the id of the synthesis method |

Continued on next page

| Option Strings | Type | Default | Help |
|---|---|---|---|
| –pcr-cycles | int | 30 | Number of cycles to be used for the PCR |
| –pcr | None | (14, None) | Specify the id of the PCR type |
| –storage-months | int | 24 | Months of storage to be simulated |
| –storage | None | (1, None) | Specify the id of the storage host |
| –sequencing | None | (2, None) | Specify the id of the sequencing method |
| –amplifier | float | 5.0 | Value by how much more distinct the errors should be |
| –probabilities | str | probabilities.json | Path to json file for error probabilities |
| –useq | str | undesired_sequences.json | Path to json file for undesired sequences |
| –gc-window | int | 50 | Size of the window to be used for the GC-Content error probability detection |
| –kmer-window | int | 10 | Size of the window to be used for the Kmer error probability detection |

**Encoder**

| Parameter | Description |
|---|---|
| rnn | Recurrent neural network, each copy of the input is encoded by seperate RNNs. |
| srnn | Recurrent neural network, returns the unencoded inputs and 1 or 2 encoded copies, depending on the rate parameter. |
| cnn | Convolutional neural network, each copy of the input is encoded by seperate CNNs. |
| scnn | Convolutional neural network, returns the unencoded inputs and 1 or 2 encoded copies, depending on the rate parameter. |
| transformer | Encoding structure based on a transformer encoder. Each copy of the input is encoded by seperate instances. |
| vae | Variational autoencoder structure. Using CNN as base neural network. |
| cnn_kernel_inc | CNN with an increasing kernel size, first pass is through a CNN with half the kernel size of the parameter value, second is with the full kernel size. |
| resnet1d | One dimensional residual neural net. |

Table 2: Implemented encoder structures.

**Transcoder**

| Parameter | Description |
|---|---|
| mlp | Multilayer perceptron, the input is split into subsequences that correspond to the number of encoded copies and transcoded by seperate MLPs. |
| cnn | Convolutional neural network. Each subsequence is encoded by seperate instances. |
| rnn | Recurrent neural network. Each subsequence is encoded by seperate instances. |
| transformer | Transcoding structure based on a transformer encoder. Each subsequence is encoded by seperate instances. |
| cnn_rnn | A combination of CNNs and RNNs. Each subsequence is encoded by seperate instances. |
| cnn_conc | A single CNN that transcodes the input sequence without splitting it into subsequences. |
| cnn_ensemble | An ensemble of CNNs, using majority voting to return the most likely candidate sequence. Only useable with binary outputs. |
| resnet | Residual neural network, the inputs are transcoded by a one dimensional ResNet together and split afterwards, to be seperatly transcoded by a linear layer. |
| resnet2d | Residual neural network, the inputs are transcoded by a two dimensional ResNet together and split afterwards, to be seperatly transcoded by a linear layer. Before the transcoding, the interleaved sequence is deinterleaved. |
| resnet2d_1d | Residual neural network for rate 1/3 only. The not interleaved inputs are transcoded by a two dimensional ResNet together, while the interleaved input is seperatly transcoded by a one dimensional ResNet. |
| resnet_ens | An ensemble of the basic ResNet component, using majority voting to return the most likely candidate sequence. Only useable with binary outputs. |
| resnet_sep | ResNet components that are inpendently trained from each other for each subsequence. |
| resnet_conc | A single, one dimensional ResNet that transcodes the input sequence without splitting it into subsequences. |
| cnn_sep | CNNs that are inpendently trained from each other for each subsequence. |

Table 3: Implemented indel reduction transcoders.

**Decoder**

| Parameter | Description |
|---|---|
| rnn | Recurrent neural network based decoder structure. |
| cnn | Convolutional neural network based decoder structure. |
| entransformer | Decoder structure based on a transformer encoder. |
| ensemble_dec | An ensemble of CNNs, using majority voting to return the most likely candidate sequence. |
| resnet1d | One dimensional residual neural net. |

Table 4: Implemented decoder structures.

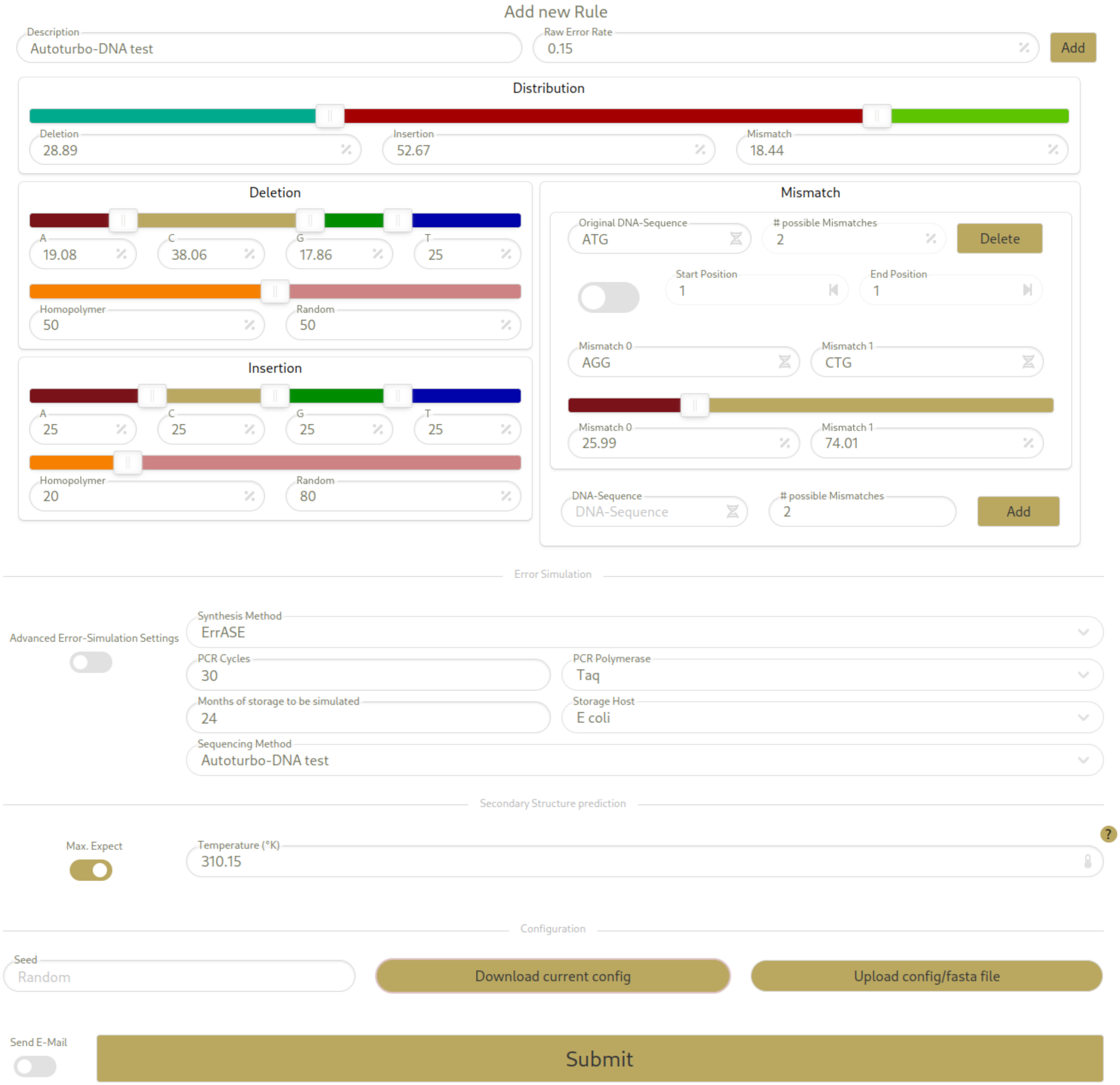


Figure 1: Example of generating a configuration file that can be used to train models with Autoturbo-DNA. Top: the MESA interface to design a new rule, showing the ability to name the rule (here ”Autoturbo-DNA test”), defining the raw error rate, the distribution of errors between deletions, insertions, and substitutions/mismatches, and the distribution and positions for each error type. For example, an insertion happens to 80 % at a random position and 20 % in a homopolymer, and the inserted base is one of the four bases with equal probability. Bottom: the simulation interface of MESA, with the new rule chosen as the sequencing method. Close to the bottom of the image, the ”Download current config” button allows the download of a JSON file containing all parameters. The JSON file data can then be used for training by adding it to the corresponding config/error_sources file of Autoturbo-DNA. The new error profile can then be selected by its ID using the related hyper-parameter (either –sequencing, –synthesis, –pcr, or –storage).

### 2 Results

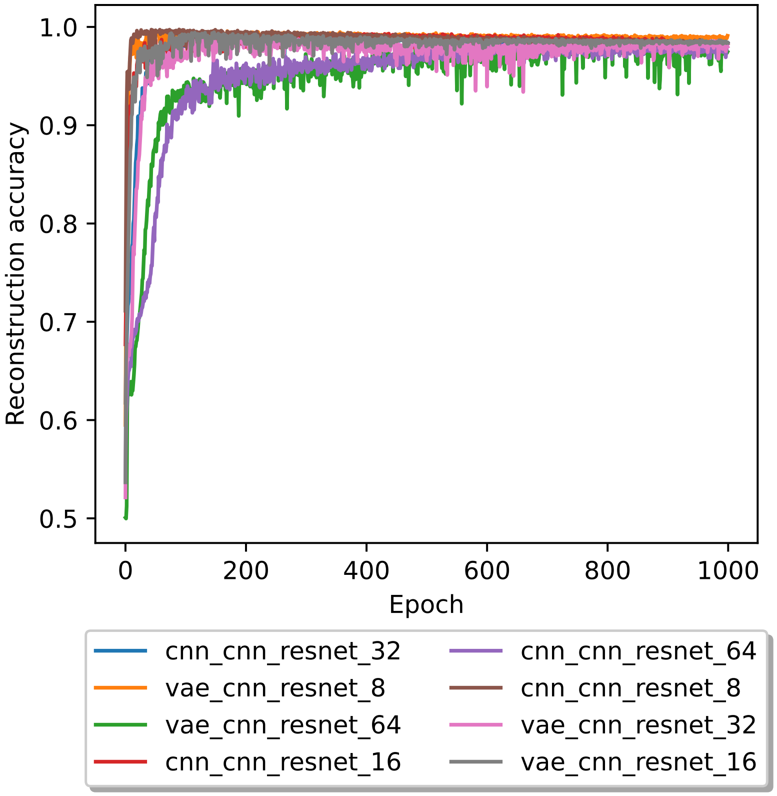


Figure 2: The reconstruction accuracy for 1000 epochs. The legend labels are structured in the form of encoder, decoder, transcoder, block size.

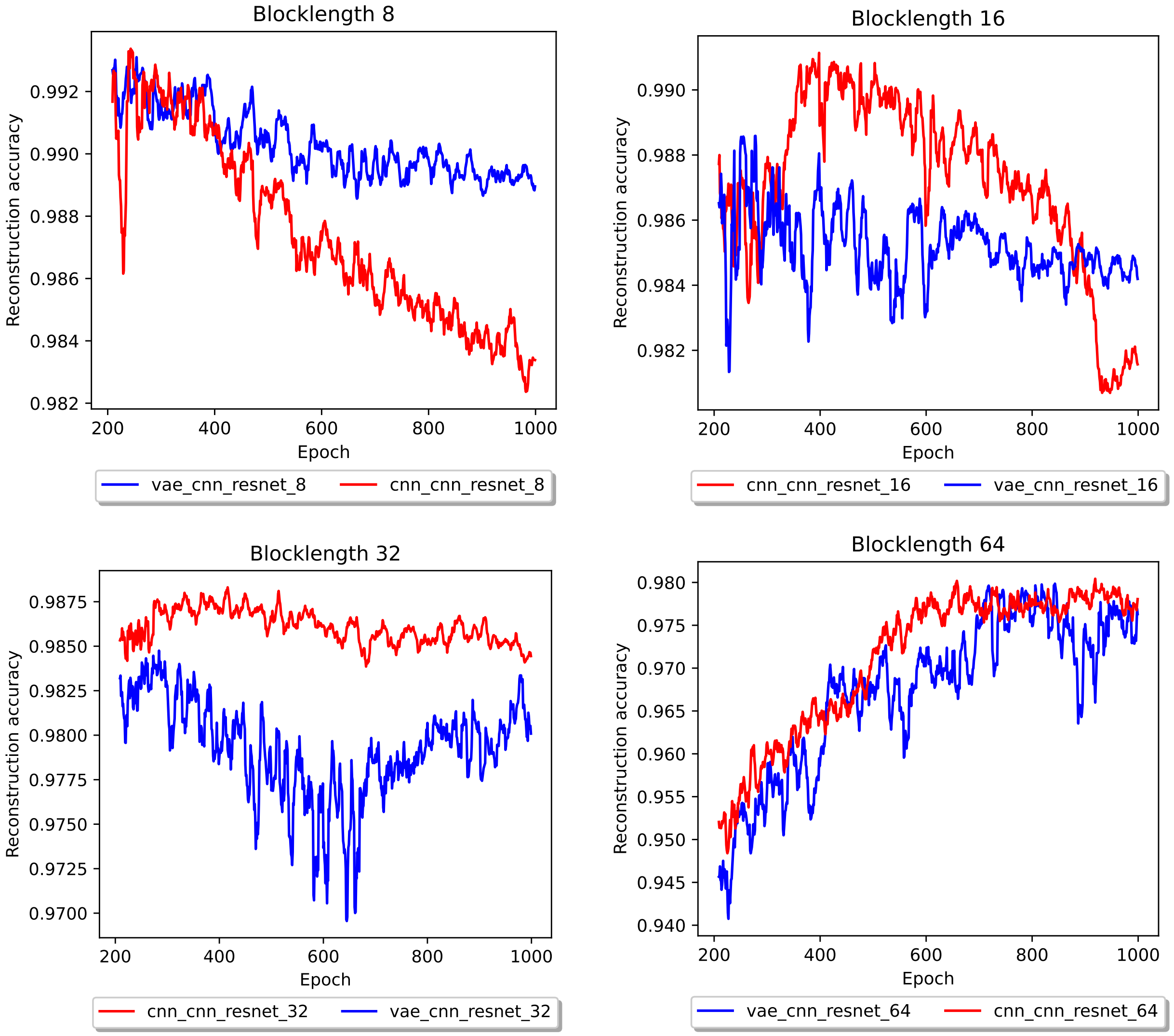


Figure 3: Reconstruction accuracy for different block lengths in a 10 epoch rolling average, beginning with epoch 200. The legend labels are structured in the form of encoder, decoder, transcoder, block size.

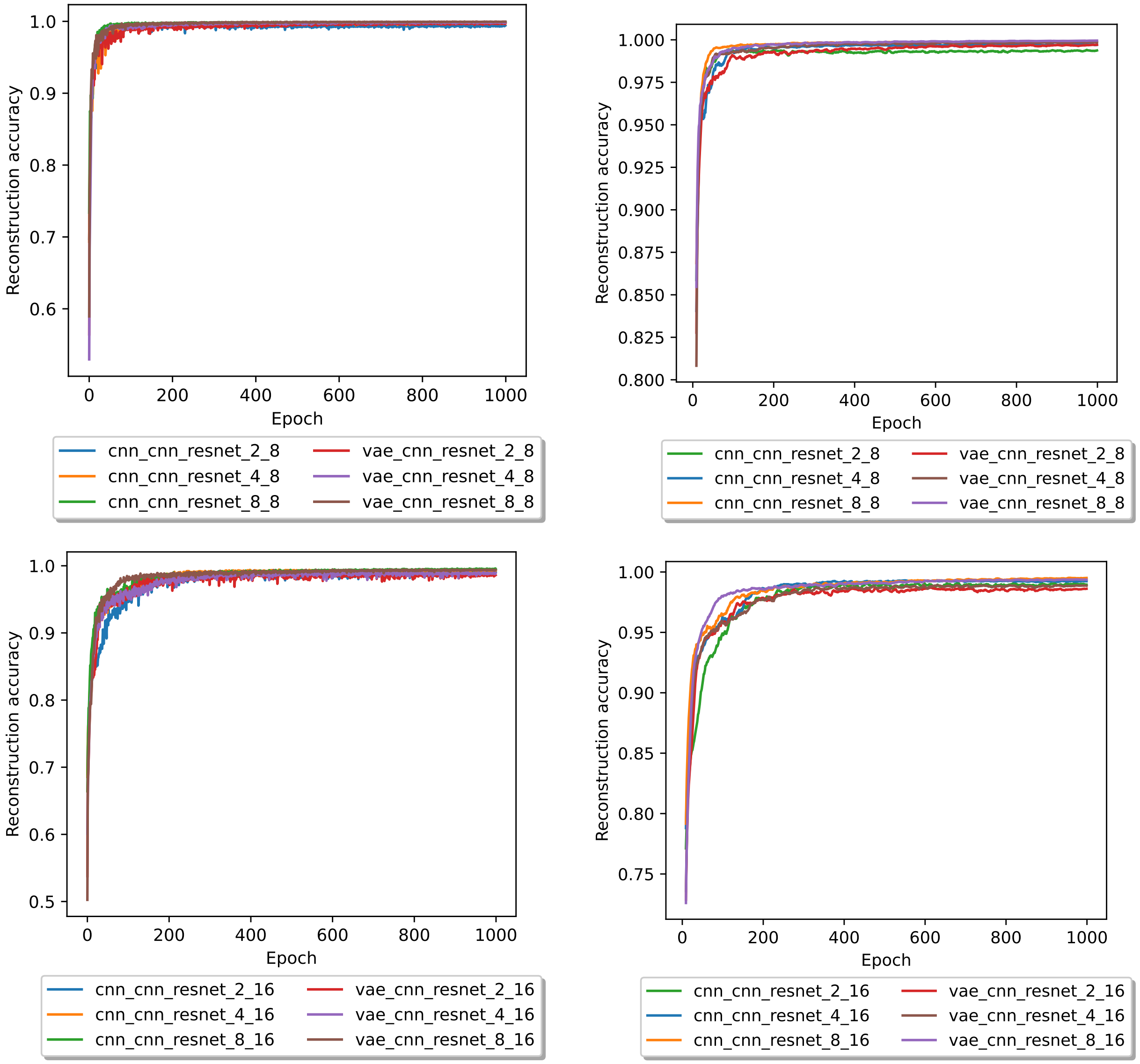


Figure 4: Reconstruction accuracy for different block lengths in a 10 epoch rolling average (right), and without the rolling average (left), for a block size of 8 (above) and 16 (below). The legend labels are structured in the form of encoder, decoder, transcoder, latent redundancy, block size.

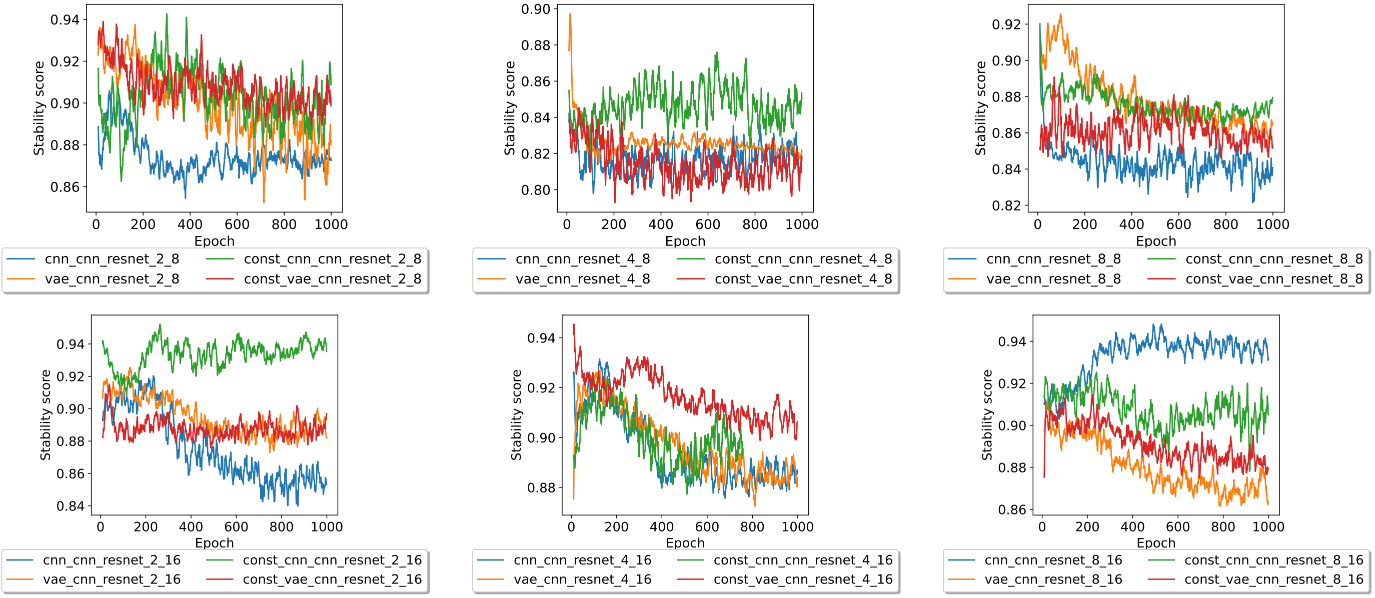


Figure 5: Stability score over 1000 epochs in a 10 epoch rolling average for a latent redundancy of 2 bits (left), 4 bits (middle) and 8 bits (right). On the top, the models were trained using a block size of 8, and on the bottom, the training was carried out with a block size of 16. The legend labels are structured in the form of constraint adherence training, encoder, decoder, transcoder, latent redundancy, block size.

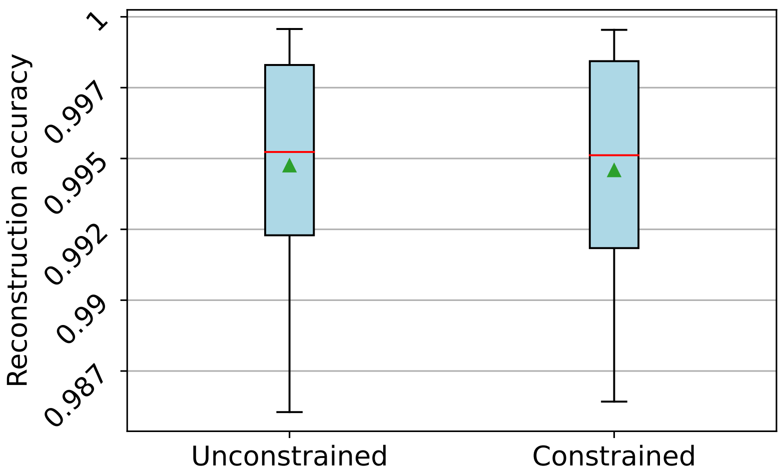


Figure 6: Boxplot of the reconstruction accuracy score of models trained without (left) and with (right) the stability score as training metric. The models were further trained with either 2, 4, or 8 bits of latent redundancy and a block size of either 8 or 16 bits. A red line represents the median, a green triangle represents the mean, and the outlier are represented by green dots.

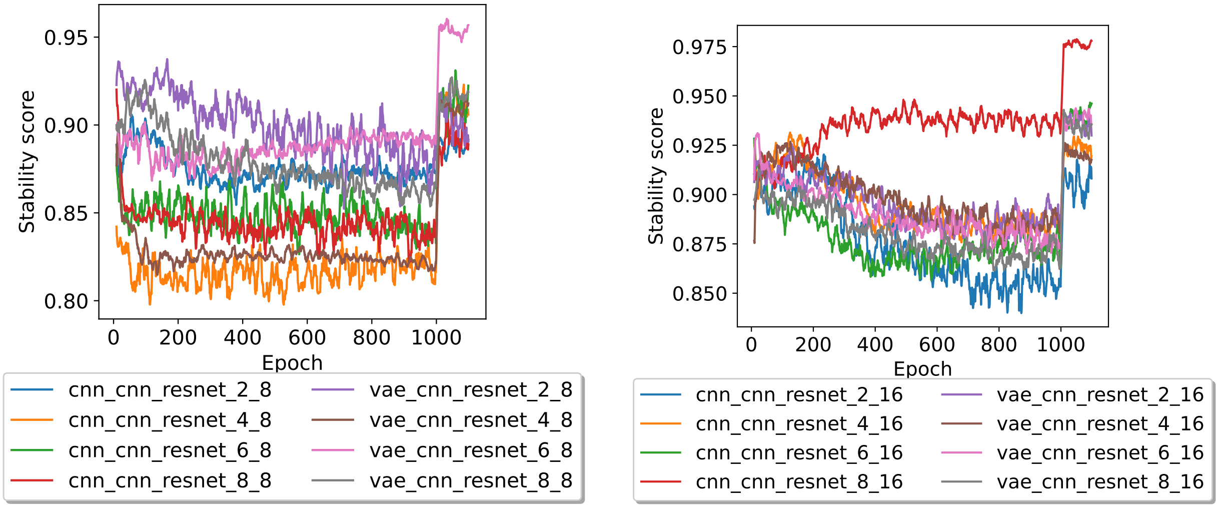


Figure 7: Stability score for different block lengths in a 10 epoch rolling average, for a block size of 8 (left) and 16 (right). Each model was trained for 1000 epochs without taking the stability score into account when training the encoder, followed by 100 epochs with the stability score being taking into account when training the encoder. The legend labels are structured in the form of encoder, decoder, transcoder, latent redundancy, block size.

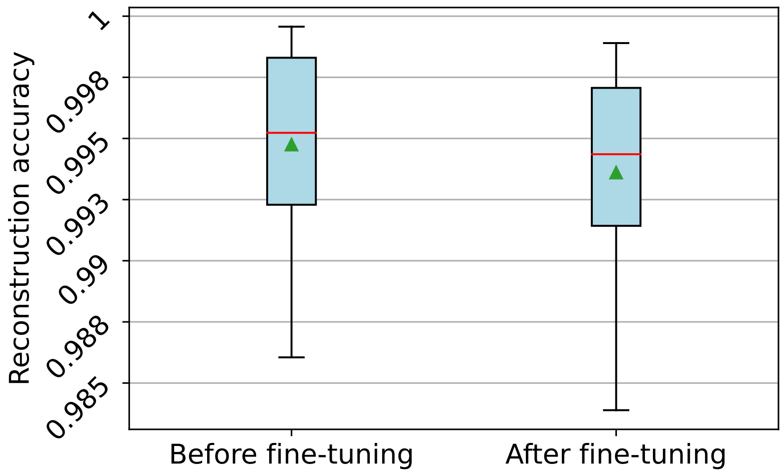


Figure 8: Boxplot of the reconstruction accuracy score of models trained before (left) and after (right) fine-tuning by utilizing the stability score as additional training metric. The models were further trained with either 2, 4, or 8 bits of latent redundancy and a block size of either 8 or 16 bits. A red line represents the median, a green triangle represents the mean, and the outlier are represented by green dots.
